# Impaired skeletal muscle regeneration induced by Cre recombinase activation in skeletal muscle stem cells

**DOI:** 10.1101/2024.01.15.575701

**Authors:** Anita Kneppers, Audrey Saugues, Carole Dabadie, Sabrina Ben Larbi, Rémi Mounier

**Affiliations:** Institut NeuroMyoGène, Physiopathologie et Génétique du Neurone et du Muscle, Université Claude Bernard Lyon 1, CNRS UMR5261, INSERM U1315, 69008 Lyon, France

## Abstract

The value of the Cre-lox system in biology is well-recognized, which is reflected by its widespread use to assess the role of a gene in a specific tissue or cell-type. Not in the least, Cre recombinase expressed under the Pax7 promotor has been invaluable for the study of skeletal muscle stem cell (MuSC) biology. In this study, we aimed to systematically assess the effects of the genetic makeup of Pax7Cre mice and Tamoxifen (Tx) treatment on skeletal muscle regeneration. We demonstrate that Tx treatment *per se* does not affect skeletal muscle regeneration at 14 days post injury (d.p.i.) induced by cardiotoxin, but specifically worsened regeneration in two Pax7Cre^ERT2^ lines. Pax7 heterozygosity in Pax7Cre^ERT2(FAN)^ mice resulted in a lower body mass and *Tibialis Anterior* (TA) mass, a higher number of fibers *per* section, and a lower number of Pax7^+^ cells than in Pax7Cre^ERT2(GAKA)^ mice, but Tx treatment did not worsen these effects caused by Pax7 haploinsufficiency. *In vitro*, proliferation of Pax7Cre^ERT2(FAN)^ MuSCs was impaired after 4-Hydrotamoxifen (4-OHT) treatment, while cell survival and differentiation remained unaffected. Together with a lower number of nuclei *per* fiber after Tx treatment in Pax7Cre^ERT2(FAN)^ and male Pax7Cre^ERT2(GAKA)^ mice, this may suggest an impaired MuSC pool expansion upon Cre activation. Yet, the *in vivo* MuSC pool was maintained in Pax7Cre mice at 14 d.p.i. Overall, our results directly show that Cre recombinase activity has an off-target effect on MuSCs, which warrants the use of Tx-treated Pax7Cre^ERT2^ mice as experimental controls in future studies, and demand caution in interpreting data using other controls in previous studies.

## INTRODUCTION

The Cre-lox system is a widely used and powerful tool in mouse genetics, allowing the spatial and/or temporal control of a target gene by deletion, insertion, translocation or inversion at specific sites in the DNA. Briefly, Cre recombinase derived from the bacteriophage P1 recombines a pair of short target sequences called *LoxP* sequences. Cre recombinase can invoke ubiquitous modification of the target gene by placing it under a promotor allowing ubiquitous expression, or can be spatially controlled invoking cell type or tissue-specific modification of the target gene by placing it under a promotor that is active in a cell-type or tissue-specific manner.

The Cre-lox system has been invaluable for the study of skeletal muscle stem cell (MuSC) biology. Notably, after identification of the essential role of the transcription factor Pax7 in MuSC specification^1^, several mouse lines were developed using Pax7 to drive MuSC-specific activation of Cre recombinase^2–4^. These lines are now widely used, and are especially valuable to study MuSC biology in adult mice due to the use of inducible Cre^ERT2^. Cre^ERT2^ encodes for a fusion protein between Cre and a modified ligand binding domain of the human estrogen receptor that interacts with heat shock protein 90 (Hsp90), retaining Cre in the cytosol. Binding of Tamoxifen (Tx) to the estrogen receptor domain releases its interaction with Hsp90, permitting entry into the nucleus, where it exerts its recombinase activity.

The use of genetically modified mice such as those expressing Pax7Cre^ERT2^ and/or a floxed allele, as well as the treatment with an exogenous estrogen-like compound such as Tx, call for appropriate experimental controls. Mademtzoglou *et al*. recently reported that nearly half of the studies that use Pax7Cre^ERT2(FAN)^ mice to specifically delete a gene of interest used wild-type mice as controls^5^. Moreover, studies using Pax7Cre^ERT2(FAN)^ mice as controls frequently do not treat these mice with Tx. However, in previous studies, we observed a differential skeletal muscle regeneration capacity between experimental controls, which may depend on both the genetic makeup and Tx treatment. For example, Mademtzoglou *et al*. reported an effect of Pax7 haploinsufficiency in Pax7Cre^ERT2(FAN)^ mice on regeneration that was worsened by Tx treatment^5^.

In this study, we aimed to systematically assess the effects of the genetic makeup of Pax7Cre mice and Tx treatment on skeletal muscle regeneration. We demonstrate that Tx treatment *per se* does not affect skeletal muscle regeneration, but worsened regeneration in two Pax7Cre^ERT2^ lines, suggesting a detrimental effect of Cre activation on MuSCs. These results warrant the use of Tx-treated Pax7Cre^ERT2^ mice as experimental controls in future studies, and demand caution in interpreting data using other controls in previous studies.

## MATERIALS AND METHODS

### Animals

Mice (male and female) were bred, housed and maintained in accordance with the French and European legislation. The experimental protocols were approved by the local ethical committee. Experiments were conducted on mice at 8-12 weeks of age, that were genotyped by PCR using toe or tail DNA. All mouse lines were maintained on a C57Bl/6 background (Fig 1). HSACre^ERT2^ mice^6^ and Pax7Cre^ERT2(FAN)^ mice^7^ were maintained heterozygous as the driver genes were (partially) replaced by the Cre^ERT2^ coding sequence. To contribute to the reduction of animals in research, Cre-negative mice from HSACre^ERT2^ and Pax7Cre^ERT2(FAN)^ breedings were used in C57Bl/6 wildtype (WT) experiments. Pax7Cre^ERT2(GAKA)^ were maintained homozygous as an internal ribosome entry site (IRES)-Cre^ERT2^ fusion protein was inserted downstream of the stop codon of PAX7.

**Figure 1.**
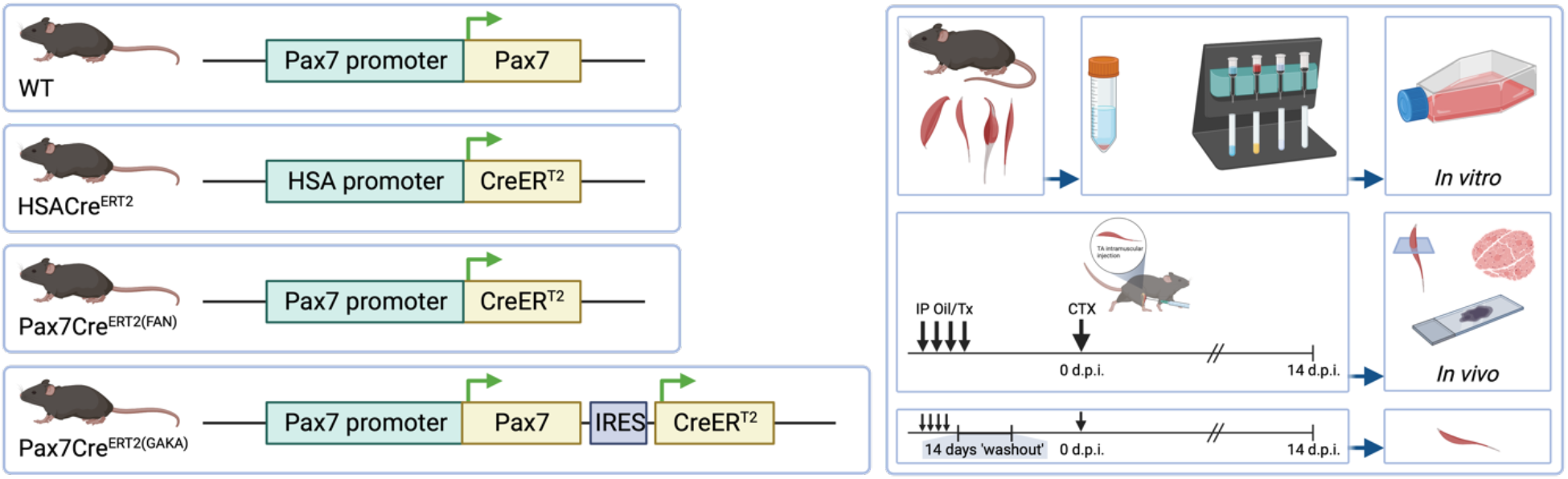
Schematic overview of mouse lines and experimental program.

Mice were treated by daily intraperitoneal injections of Tamoxifen (Tx; 0.1mg/g body weight) or an equal volume of vehicle (Sunflower Oil) for 4 days. Subsequent skeletal muscle injury was induced by Cardiotoxin (CTX) injection in the *Tibialis Anterior* (TA) (50μl *per* TA, 12μM). Skeletal muscle injury was induced 1 week after the first Tx or Oil injection, or 3 weeks after the first injection in experiments with an additional 14 days ‘washout’ (Fig 1).

### Primary MuSC isolation and culture

MuSCs were isolated from hindlimb muscles of untreated WT, HSACre^ERT2^, Pax7Cre^ERT2(FAN)^, or Pax7Cre^ERT2(GAKA)^ male mice, as described before^8^. Briefly, muscles were dissected and digested with Collagenase II and Dispase. Dissociated muscles were passed through a 70μm filter, and red blood cells were lysed in ACK lysis buffer. Cell suspensions were then incubated with magnetic beads directed to non-target cells, and run through a MACS Column in the magnetic field of a MACS Separator to retain non-target cells. Flow through, enriched for MuSCs was seeded onto Gelatin coated supports at 3000 cells/cm^2^, and amplified in proliferation medium (DMEM F12, 20% Foetal Bovine Serum (FBS), 2% Ultroser G, and 1% Penicillin/Streptomycin (PS)). MuSC purity after MACS was verified by Desmin immunohistochemistry (IHC) to be >97%.

After amplification, MuSC progeny were trypsinized and plated at indicated densities onto Gelatin coated supports (when kept in proliferating conditions) or onto Matrigel coated supports (when switched to differentiation conditions^8^). To induce differentiation, proliferation medium was removed and replaced by differentiation medium (DMEM F12, 2% Horse Serum (HS), 1% PS). To assess proliferation, cells were incubated with 5 ethynyl 2’ deoxyuridine (EdU; 10μM).

### Immunohistochemistry

For immunohistochemical analyses, TA muscles were isolated, embedded in tragacanth gum, frozen in liquid nitrogen cooled isopentane, and stored at -80°C until use. 10 μm-thick cryosections were prepared for hematoxylin-eosin (HE) staining and immunolabeling. HE staining was used to assess the efficiency of CTX injections, and muscles were only kept for further analyses if ≥75% of the muscle area consisted of centrally nucleated fibers.

For immunolabeling, cryosections were permeabilized for 10 minutes (min) in 0.5% Triton X-100 in PBS and saturated in 2% BSA for 1 hour (hr) at room temperature (RT). For the detection of myofibers, sections were labelled overnight at 4°C with primary antibodies directed against Laminin α1 (1:200). For identification of MuSCs, sections were fixed for 20 min in 4% paraformaldehyde (PFA) at RT, and permeabilized for 6 min in 100% methanol at -20°C. After antigen retrieval for 2*5 min in 10 mM Citrate buffer at 90°C, sections were saturated in 4% BSA for 2 hr at RT. Sections were then co-labelled overnight at 4°C with primary antibodies directed against Paired Box 7 (Pax7; 1:50) and Laminin α1 (1:100) in 2% BSA. Secondary antibodies were coupled to FITC, Cy3 or Cy5 (Jackson ImmunoResearch Inc.; 1:200) and incubated for 45-60 min at 37°C. Nuclear counterstain was performed by 10 seconds incubation with Hoechst (2 μM), and coverslips were mounted with Fluoromount G.

Cultured cells were fixed for 10 min in 4% PFA at RT, permeabilized for 10 min in 0.5% Triton X-100 in PBS, and saturated in 4% BSA for 1 hr at room temperature (RT). Cells were then incubated overnight at 4°C with primary antibodies directed against Desmin (1:200) and Myogenin (MyoG; 1:50) in 2% BSA. Secondary antibodies were coupled to FITC, Cy3 or Cy5 (Jackson ImmunoResearch Inc.; 1:200) and incubated for 45-60 min at 37°C. Labelling of EdU was performed after secondary antibody incubation using Click-It EdU Kit. Nuclear counterstain was performed by 10 sec incubation with Hoechst, and cells were covered with Fluoromount G.

Images of fluorescent immunolabeling in cell culture supports and scanned slides were acquired with a Zeiss Axio Observer Z1 connected to a Coolsnap HQ2 camera. For each condition of each experiment, either full sections, or at least 5-10 randomly chosen fields of view were counted. Fiber cross-sectional area and fiber number were determined on whole sections using the Open-CSAM ImageJ macro^9^.

### Flow cytometry

For flow cytometric analyses, cultured MuSCs were detached by trypsinization after 24 hours of 4-Hydroxytamoxifen (4-OHT; 500nM) or vehicle (EtOH) treatment. For cell cycle assessment, MuSCs were incubated with EdU during the last hour before trypsinization. Cells were then subjected to staining with Click-iT EdU, Ghost Dye, and ɣ-H2AX antibody. Briefly, cells in suspension were first incubated with Ghost Dye for 30 min at 4°C. Cells were then fixed in 4% PFA for 15 min at RT, permeabilized in 0.5% Triton X-100 for 10 min at 4°C, and stained with ɣ-H2AX antibody for 20 min at 4°C. Finally, EdU was detected by Click-iT reaction for 30 min at RT. After washing, cells were stained with DAPI, and analysed using a flow cytometer (LSRII, BD).

### Statistical analyses

Bars represent means +/- SEM. For *in vitro* experiments, paired replicates signify cells extracted from individual mice. For *in vivo* experiments, replicates signify individual muscles. Normal distribution was approximated from QQ plots. All *in vivo* results were analysed using unpaired parametric analyses, whereas *in vitro* results were analysed using paired parametric analyses. Statistical significance of Tx treatment effects was determined using two-sided Student’s *t* tests or Welch’s t tests, and significance of genotype or genotype*Tx treatment effects was determined by 2-way ANOVA, using GraphPad Prism software version 10.0.3.

### Key resources

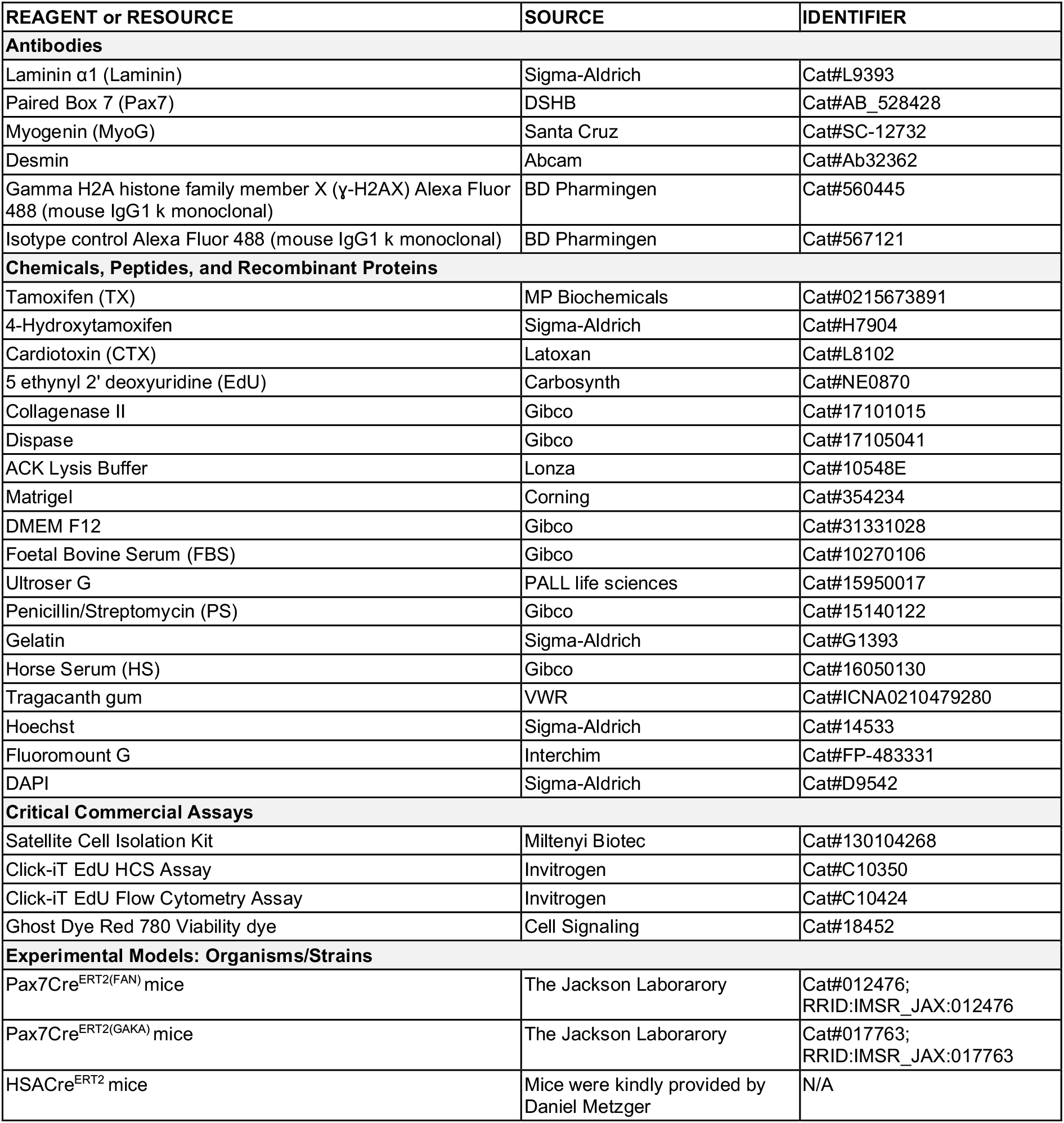

## RESULTS

### Tx treatment does not impair skeletal muscle regeneration after injury in C57Bl/6 mice

Estrogens and estrogen-like compounds are known modulators of skeletal muscle regeneration, as reviewed before ^e.g., 10,11^. We therefore asked if Tx treatment *per se* could cause alterations in skeletal muscle regeneration. To this end, we treated C57Bl/6 male and female mice for 4 days with intraperitoneal Tx or vehicle (Oil), and subjected *Tibialis Anterior* (TA) muscles to cardiotoxin (CTX)-induced injury at 7 days after the first Tx injection.

At 14 days post injury (d.p.i.), neither body mass nor absolute TA mass were affected in Tx-treated male mice (Fig 2A-B), while relative TA mass even tended to be higher after Tx treatment (Fig 2C). Furthermore, the average myofiber cross-sectional area (CSA) remained unaltered in males (Fig 2D). Similarly, body mass was unaffected in Tx-treated female mice (Fig 2E). In female mice, TA mass of female mice even tended to be higher after Tx treatment (Fig 2F), but was unaffected when corrected for body mass (Fig 2G). Furthermore, CSA also remained unaffected in females (Fig 2H), together demonstrating that Tx treatment *per se* does not impair skeletal muscle regeneration at 14 d.p.i.

**Figure 2.**
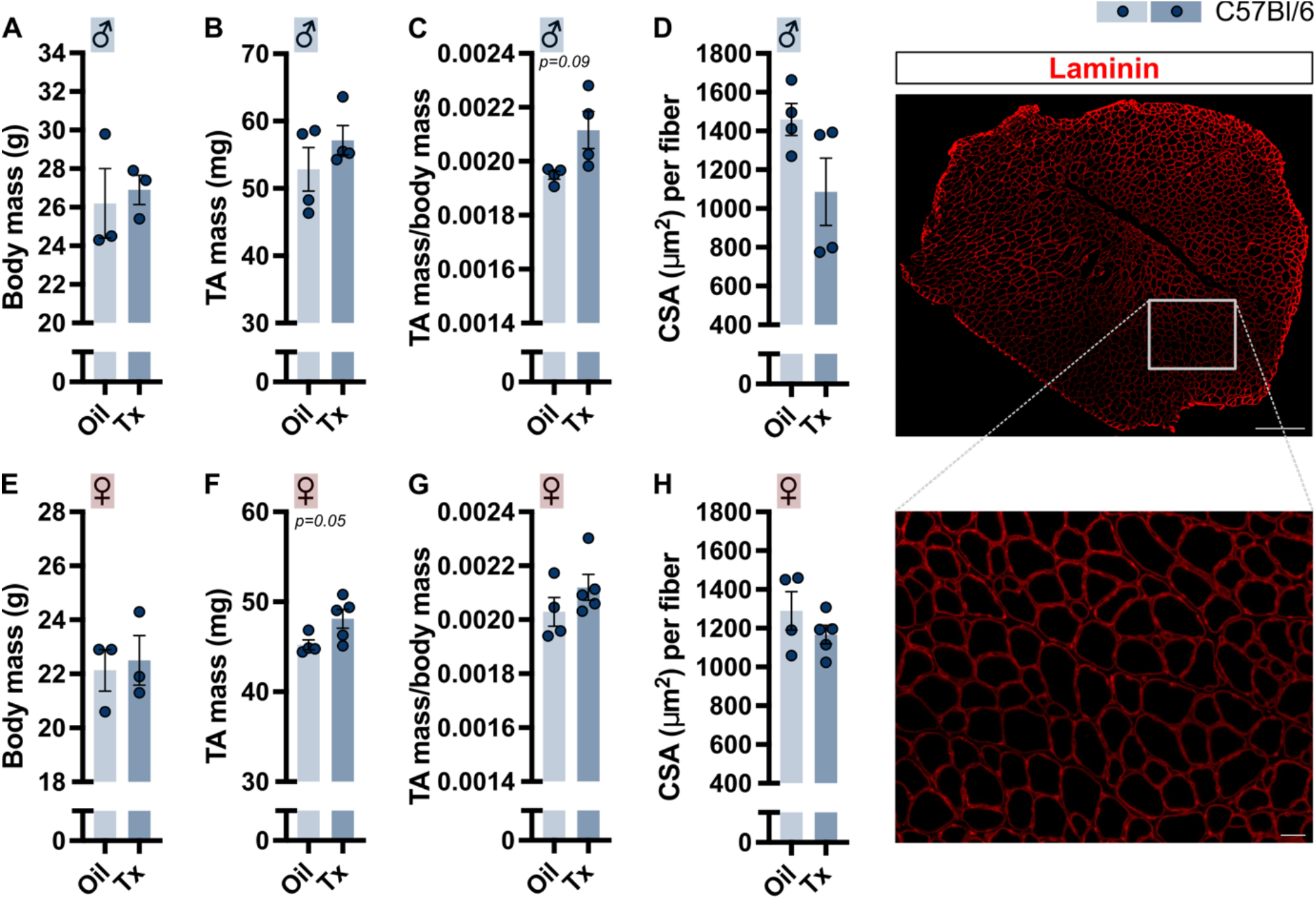
Skeletal muscle regeneration after Tamoxifen treatment at 14 days post injury in C57Bl/6 mice. Male mice treated with Oil or Tamoxifen (Tx) (A-D). A) Body mass, B) *Tibialis Anterior* (TA) mass, C) TA mass *per* mg body mass, D) average fiber cross sectional area (CSA) (left) and represented image of TA section stained for Laminin (right). Female mice treated with Oil or Tx (E-H). E) Body mass, F) TA mass, G) TA mass *per* mg body mass, H) average fiber CSA. Bars represent means +/- SEM.

### Tx treatment specifically impairs skeletal muscle regeneration after injury in Pax7Cre^*ERT2*^ *mice*

To assess if Tx treatment has an effect on skeletal muscle regeneration in Pax7Cre^ERT2^ mice, we subjected both mice harboring the FAN allele and the GAKA allele to Tx treatment and subsequent CTX-induced injury.

At 14 d.p.i., body mass of male Pax7Cre^ERT2^ mice was unaffected by Tx treatment (Fig 3A-B). However, relative TA mass and average fiber CSA at 14 d.p.i. were lower upon Tx treatment in male FAN mice (Fig 3C-D). Similarly, absolute and relative TA mass, as well as average fiber CSA tended to be lower at 14 d.p.i. after Tx treatment in male GAKA mice (Fig 3B-D), overall demonstrating a detrimental effect of Tx treatment on skeletal muscle regeneration in Pax7Cre^ERT2^ mice. This detrimental effect did not seem to depend on sex. Indeed, body mass of female Pax7Cre^ERT2^ mice was unaffected by Tx treatment at 14 d.p.i. (Fig 3E). Furthermore, female FAN mice tended to have a lower absolute and relative TA mass, as had a lower average fiber CSA at 14 d.p.i. after Tx treatment (Fig 3F-H). Both absolute and relative TA mass were lower in female GAKA mice, yet their average fiber CSA remained unaffected by Tx treatment at 14 d.p.i. (Fig 3F-H). Importantly, Tx treatment responses did not differ significantly between FAN and GAKA mice for any of the parameters, but both male and female GAKA mice had a higher body mass (Fig 3A&E). Furthermore, male GAKA mice had a lower relative TA mass than FAN mice (Fig 3C), while female GAKA mice had a higher absolute TA mass than FAN mice that did not result in a difference in relative TA mass (Fig 3F-G).

**Figure 3.**
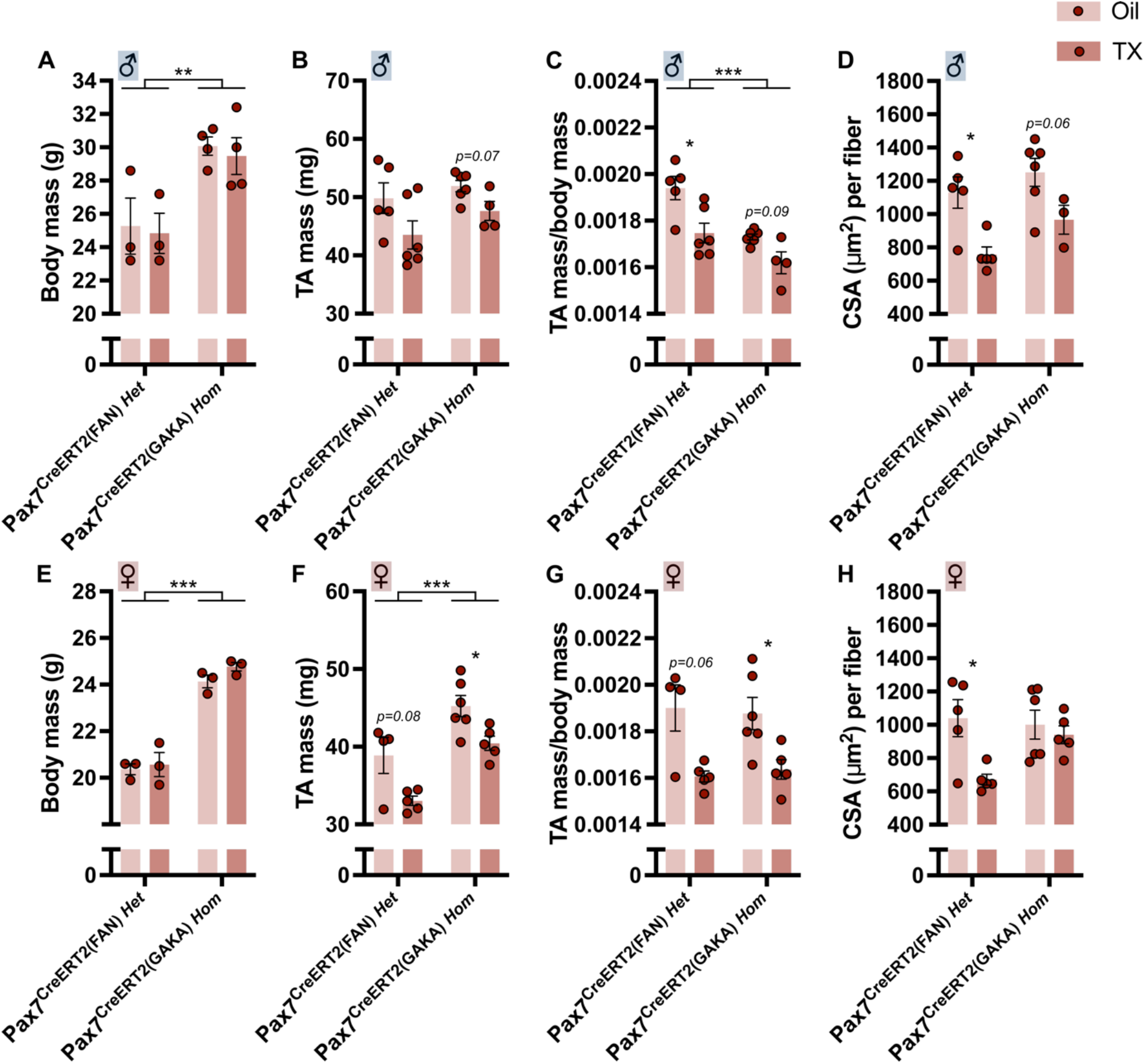
Skeletal muscle regeneration after Tamoxifen treatment at 14 days post injury in Pax7Cre^ERT2^ mice. Male mice treated with Oil or Tamoxifen (Tx) (A-D). A) Body mass, B) *Tibialis Anterior* (TA) mass, C) TA mass *per* mg body mass, D) average fiber cross sectional area (CSA) Female mice treated with Oil or Tx (E-H). E) Body mass, F) TA mass, G) TA mass *per* mg body mass, H) average fiber CSA. Bars represent means +/- SEM. *p<0.05, **p<0.01 and ***p<0.001.

The observed detrimental effects of Tx treatment on skeletal muscle regeneration might be due to the activation of Cre. Nevertheless, Tx treatment in male HSACre^ERT2^ mice does not affect body mass, absolute and relative TA mass, or average fiber CSA at 14 d.p.i. (Fig S1A-D). Interestingly, absolute and relative TA mass, as well as average fiber CSA were lower at 14 d.p.i. upon Tx treatment in female HSACre^ERT2^ mice (Fig S1E-H). While striking, the gender effect of Tx treatment in HSACre^ERT2^ mice will not be further dissected in this study.

Overall, this demonstrates that Tx treatment specifically impairs skeletal muscle regeneration in both male and female Pax7Cre^ERT2^ mice, and suggests a specific detrimental effect of Cre activation in MuSCs. To assess if such effects persist after cessation of Tx-induced Cre activation, male Pax7Cre^ERT2^ mice harboring the FAN allele were subjected to the same Tx treatment, but muscle injury was induced by CTX injection with an additional 14 days ‘washout’ period (*i*.*e*., 21 days after the first Tx injection, Fig 1). At 14 d.p.i., body mass of these mice was unaffected (Fig S1I), and similar to our previous study, absolute and relative TA mass remained lower at 14 d.p.i. upon Tx treatment (Fig S1J-K). Moreover, Tx treatment responses did not statistically differ between male FAN mice with or without 14 days of delay before CTX-induced skeletal muscle injury, together suggesting that Cre activity and/or its detrimental effects on MuSCs persist.

### Tx treatment specifically reduces the fusion index at 14 days post injury in Pax7Cre^*ERT2*^ *mice*

Fiber characteristics were further explored to address the cellular cause of the Tx-induced lower TA mass and fiber CSA at 14 d.p.i. The number of fibers *per* section tended to be increased upon Tx treatment in male Pax7Cre^ERT2^ mice harboring the FAN allele, but was unaffected in male Pax7Cre^ERT2^ mice harboring the GAKA allele at 14 d.p.i. (Fig 4A), resulting in a statistically different Tx response between male FAN and GAKA mice (p=0.039). Importantly, both male FAN and GAKA mice had a lower number of nuclei *per* fiber (Fig 4B). Similarly, the number of fibers *per* section was unaffected by Tx treatment in female FAN and GAKA mice at 14 d.p.i. (Fig 4C), while FAN but not GAKA female mice had a lower number of nuclei *per* fiber (Fig 4D). The Tx-induced reduction in number of nuclei *per* fiber did not statistically differ between FAN and GAKA mice (Fig 4B&D). Notably, the number of fibers *per* section at 14 d.p.i. was lower in both male and female GAKA mice than in FAN mice, while the number of nuclei *per* fiber was similar between both genotypes (Fig 4A-D).

**Figure 4.**
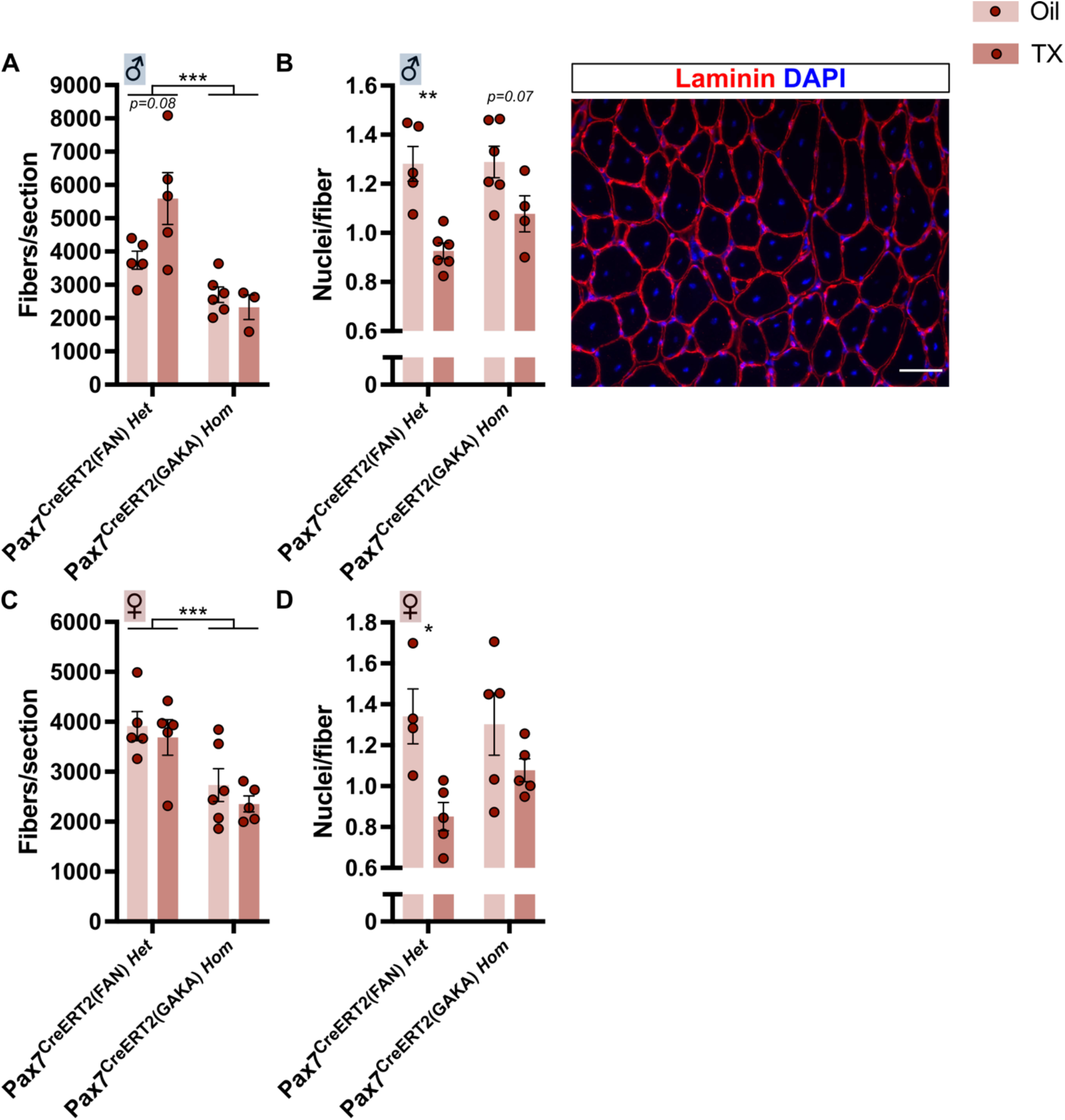
Myofiber characteristics after Tamoxifen treatment at 14 days post injury in Pax7Cre^ERT2^ mice. Male mice treated with Oil or Tamoxifen (Tx) (A-B). A) Number of fibers *per* section, B) Number of nuclei *per* fiber (left) and representative image (right). Female mice treated with Oil or Tx (C-D). C) Number of fibers *per* section, D) Number of nuclei *per* fiber. Bars represent means +/- SEM. *p<0.05, **p<0.01 and ***p<0.001.

In line with the unaffected skeletal muscle regeneration upon Tx treatment in C57Bl/6 mice, the number of fibers *per* section was unaltered in both male and female C57Bl/6 mice at 14 d.p.i. (Fig S2A-B). Similarly, the number of nuclei *per* fiber was unaltered upon Tx treatment in both male and female HSACre^ERT2^ mice (Fig S2C-D).

Altogether, this shows that Tx treatment specifically reduces the fusion index after injury in Pax7Cre^ERT2^ mice, and suggests an impaired MuSC contribution to *de novo* myofiber formation upon MuSC-specific Cre activation.

### Cre activation in MuSCs impairs MuSC pool expansion

Since MuSCs are the regenerative precursors for *de novo* myofiber formation after injury, their function upon Tx treatment was assessed. Tx treatment did not affect the MuSC pool at 14 d.p.i. in either FAN or GAKA Pax7Cre^ERT2^ mice, independent of sex (Fig 5A-B). However, specifically in females, the number of MuSCs *per* fiber was higher in GAKA than in FAN mice (Fig 5B). Similar to Pax7Cre^ERT2^ mice, the MuSC pool was unaffected by Tx treatment in male and female C57Bl/6 mice (Fig S3A-B), and in male HSACre^ERT2^ mice (Fig S3C). Surprisingly, Tx treatment tended to reduce the MuSC pool at 14 d.p.i. specifically in female HSACre^ERT2^ mice (Fig S3D). Together, this indicates that neither Tx treatment *per se*, nor Cre activation in MuSCs alters MuSC pool maintenance at 14 d.p.i. To further dissect the effect of Cre activation on MuSC function, MuSCs were isolated from male Pax7Cre^ERT2^ mice harboring the FAN allele and expanded *in vitro*. Treatment of FAN MuSCs for 24 hours with the active Tx metabolite 4-OHT reduced proliferation by 13% (Fig 5C-D). In line, the fraction of cells in S-phase tended to be lower after 24 hours of 4-OHT treatment (Fig 5E). Cell survival was unaffected by 24 hours of 4-OHT treatment (Fig 5F). Moreover, the reduced proliferation capacity does not seem to be due to a precautious cell cycle exit, evidenced by the unaltered fraction of differentiated cells after 24 hours of 4-OHT treatment in proliferating conditions (Fig 5G). Finally, also the MuSC intrinsic differentiation capacity was unaffected by 24 hours of 4-OHT treatment in differentiating conditions (Fig 5H). Together, these data indicate that Cre activation in MuSCs affects MuSC pool expansion.

**Figure 5.**
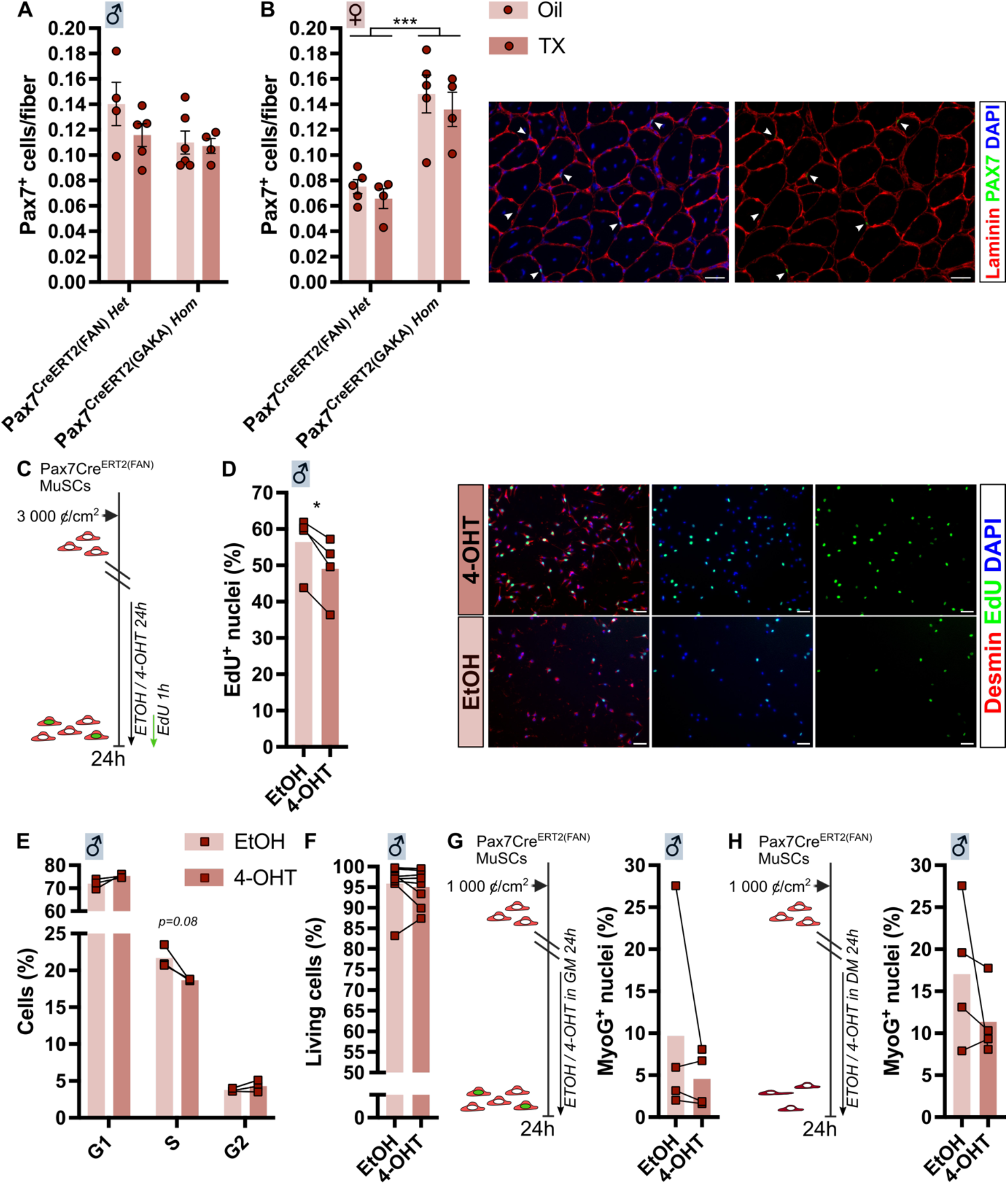
MuSC number and function after Cre activation in Pax7Cre^ERT2^ MuSC. A) number of Pax7^+^ MuSCs *per* fiber at 14 days post injury (d.p.i.) in male Pax7Cre^ERT2^ mice, B) number of Pax7^+^ MuSCs *per* fiber at 14 d.p.i. in female Pax7Cre^ERT2^ mice, C) schematic of *in vitro* experiments on cells derived from male Pax7Cre^ERT2(FAN)^ mice in panel D-G, D) percentage of EdU^+^ nuclei in myogenic cells (desmin^+^) by immunofluorescence (IF), E) cell cycle partition by flow cytometry, F) percentage of living cells by flow cytometry, G) percentage of Myogenin (MyoG) expressing nuclei in myogenic cells in proliferation medium, H) schematic of experimental setup (left) and percentage of MyoG expressing nuclei in myogenic cells in differentiation medium (right). Bars represent means +/- SEM. *p<0.05 and ***p<0.001.

### Cre off-target effects in MuSCs and beyond

Although Pax7Cre^ERT2^ mice have been proven crucial to study MuSC biology, and are now widely used in the field, a potential off-target effect of Cre activity in these driver mice has only been recently suggested^5^. Notably, this study mainly ascribes observed Tx-induced effects to the genetic makeup of Pax7Cre^ERT2^ mice harboring the FAN allele, which are heterozygous for a Pax7 knockout allele resulting in Pax7 haploinsufficiency. However, several previous studies have reported direct off-target effects of Cre activity *in vivo* in mice (Table 2). Generally, these studies observe impaired cell proliferation/pool expansion, and nearly all observe signs of apoptosis in cells exposed to active Cre recombinase. Moreover, various studies describe chromosomal aberrations and a DNA damage response upon exposure to active Cre recombinase. To assess if Cre activation induces a DNA damage response in MuSCs, 4-OHT treated MuSCs were stained for ɣ-H2AX and assessed by flow cytometry. We observed no alterations in the median fluorescent intensity of ɣ -H2AX in either phase of the cell cycle (Fig S3E-F). Interestingly, 4-OHT treatment tended to increase cell size (Fig S3G), which may point to an induction of senescence.

**Table 2.**
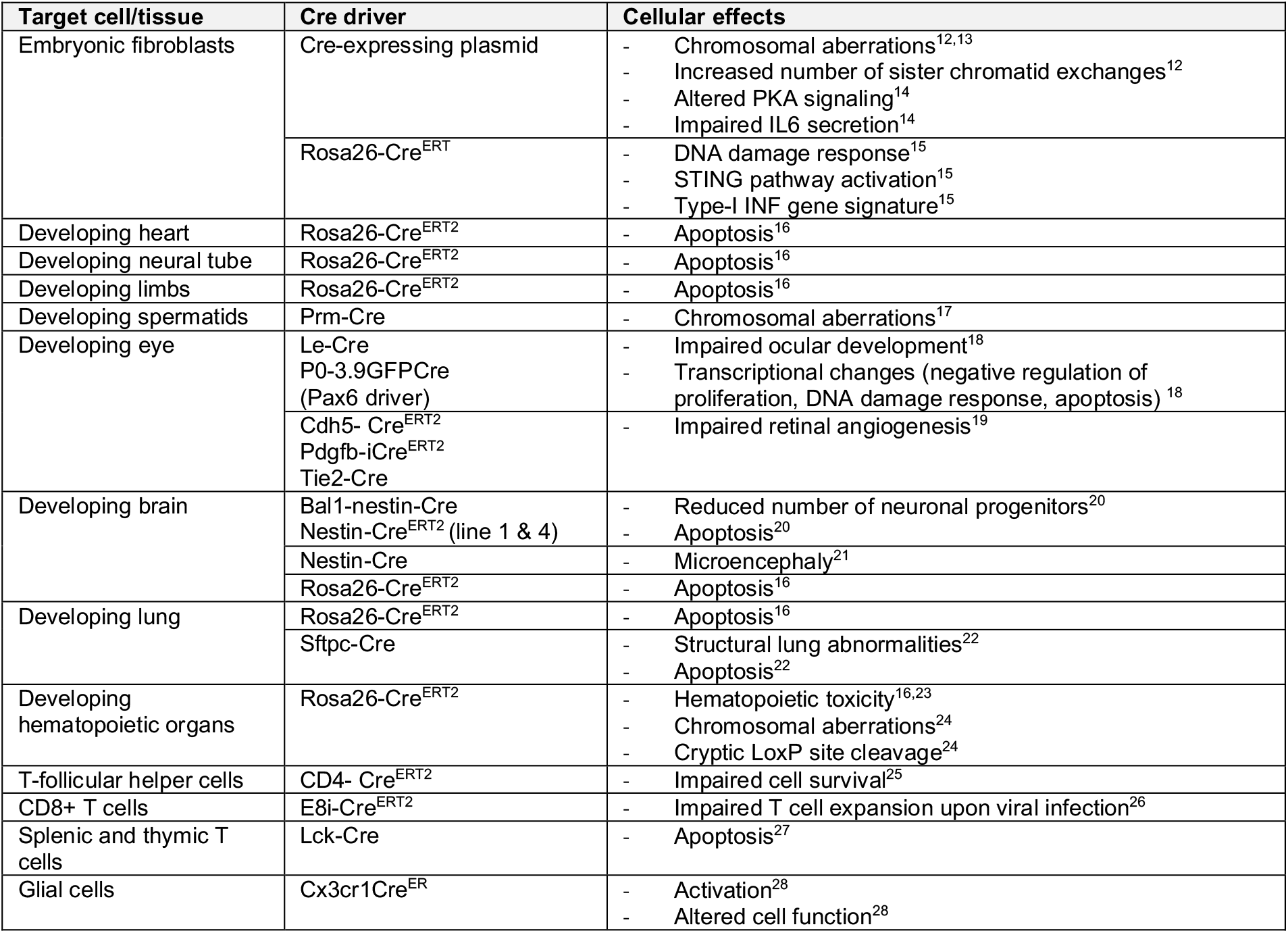
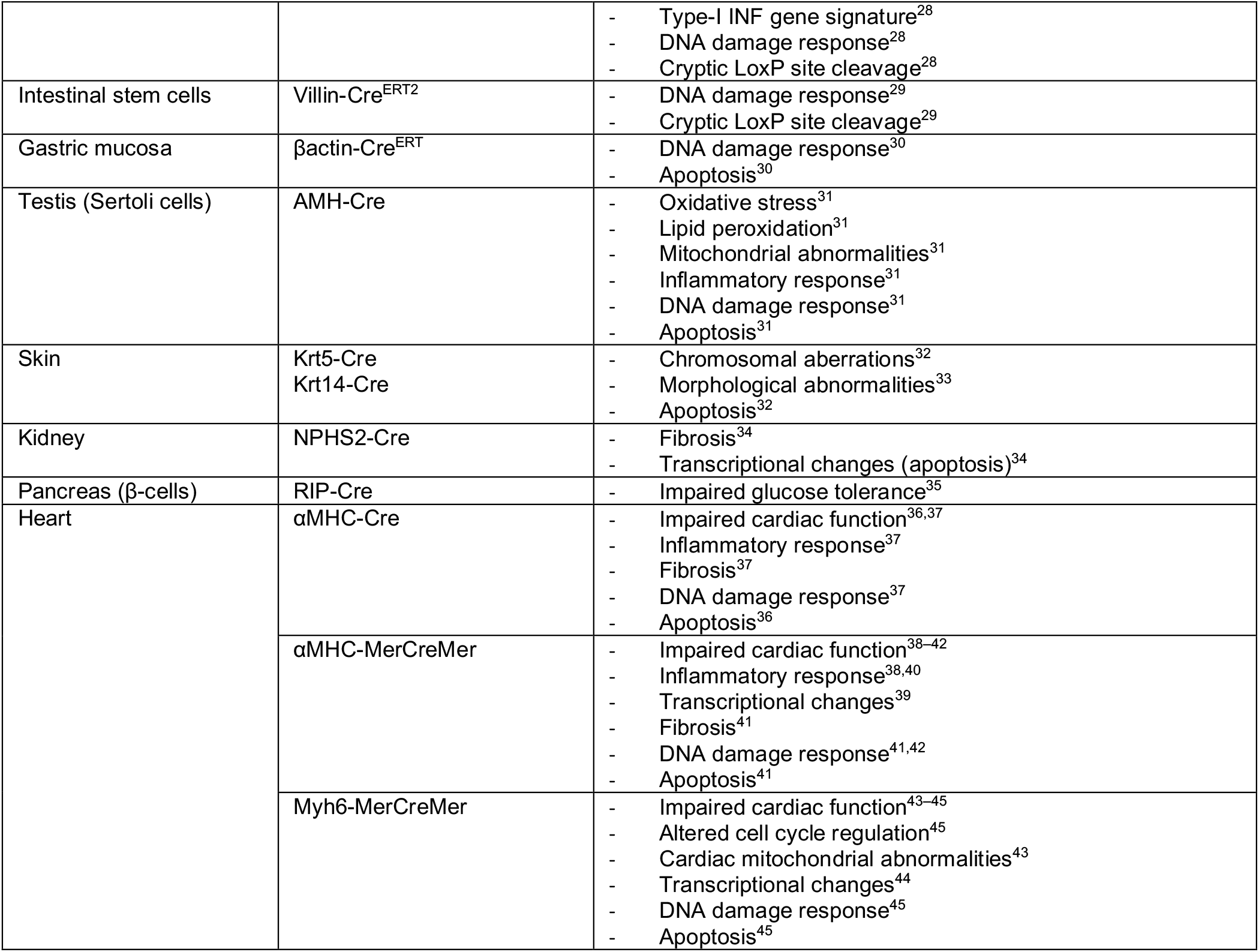
Reported Cre off-target effects *in vivo* in mice.

## DISCUSSION

The value of the Cre-lox system in biology is well-recognized, which is reflected by its widespread use to assess the role of a gene in a specific tissue or cell-type. Not in the least, Cre recombinase expressed under the Pax7 promotor has been invaluable for the study of MuSC biology. Yet, Pax7Cre^ERT2^ mice come with some caveats, that are known within the field but often not empirically demonstrated or reported.

Mademtzoglou *et al*. recently demonstrated an impaired skeletal muscle regeneration in Pax7Cre^ERT2(FAN)/+^ mice due to Pax7 haploinsufficiency caused by Pax7 heterozygosity^5^. They observed these effects under prolonged Tx treatment, and in agreement with the laboratory that generated these mice, recommend the use of Tx treated Pax7Cre^ERT2(FAN)/+^ controls. In our study, Pax7 heterozygosity in Pax7Cre^ERT2(FAN)^ mice resulted in a lower body mass and TA mass, a higher number of fibers *per* section, and a lower number of Pax7 cells than in Pax7Cre^ERT2(GAKA)^ mice. However, Tx treatment did not worsen these effects caused by Pax7 haploinsufficiency. Importantly, we demonstrate that Tx treatment impaired skeletal muscle regeneration in both Pax7Cre^ERT2(FAN)^ and Pax7Cre^ERT2(GAKA)^ mice, but not in WT or male HSACre^ERT2^ mice, strongly suggesting a detrimental effect of Cre recombinase activity in MuSCs.

Numerous studies revealed a toxicity of Cre activity in other tissues and cell types (Table 1), most observing impaired cell proliferation/pool expansion, and nearly all observe signs of apoptosis in cells exposed to active Cre recombinase. Moreover, many studies describe chromosomal aberrations and a DNA damage response upon exposure to active Cre recombinase. Although we demonstrate evidence of impaired MuSC pool expansion upon Cre activation, our analyses did not reveal an increase in MuSC death or a DNA damage response after 24 hours of Cre activation *in vitro*. However, this does not exclude a potential contribution of DNA damage or cell death to impaired MuSC pool expansion. Notably, Pax7Cre^ERT2(FAN)^ MuSC were larger after Cre activation *in vitro*, which is a sign of senescence, that could be induced by prior DNA damage or chromosomal aberrations^46^. Overall, the direct consequence of Cre activation in MuSCs remains unknown.

Interestingly, the mammalian genome contains cryptic LoxP sites that can be targeted by Cre recombinase^47^, and three studies demonstrate cryptic LoxP site cleavage directly linking it to their Cre off-target effects^24,28,29^. Perhaps, Cre-induced chromosomal alterations and cryptic LoxP site cleavage are good first targets for future exploration in MuSCs. It would also be interesting to assess if such ‘illegitimate’ chromosomal cuts at cryptic LoxP sites occur even in the presence of introduced LoxP sites.

Given the value of Pax7Cre^ERT2^ in MuSC research, it would be essential to define the condition that minimizes Cre off-target effects. We aimed to avoid Cre off-target effects by addition of a 14-day washout period before CTX-induced injury, but did not recover the impaired regenerative capacity after Tx treatment. This may suggest that the off-target effects of Cre activity are permanent, and are only recovered after (competitive) removal of damaged cells. Alternatively, the fat-soluble Tx may be stored in tissues, inadvertently prolonging the exposure of target cells to Cre activity. Several studies link Cre off-target effects to the level of Cre expression and activity^20,25,36,39,41,44^. As such, it seems reasonable to optimize the Tx dose and duration to avoid Cre off-target effects while maintaining sufficient recombination of the target gene – an effort that should be repeated for each new floxed allele. It should also be noted that we used homozygous Pax7Cre^ERT2(GAKA)^ mice in the current study. Perhaps, a lower Cre expression in heterozygous Pax7Cre^ERT2(GAKA)^ mice could (partially) avoid the detrimental effect of Cre recombinase activity in MuSCs.

An interesting development is the generation of self-deleting Cre constructs, which can reduce Cre toxicity^13,48,49^. However, their capacity to retain sufficient recombination of the target gene should be validated, as the recombination of one allele does not always faithfully ‘report’ recombination of another allele^50^.

Despite the cautionary tales from other fields – all concluding the need for controls with active Cre recombinase (all references in Table 1), the literature inventory by Mademtzoglou *et al*. demonstrates that the use of controls without Cre expression or activation is still common practice in MuSC research^5^. Our results directly show that Cre recombinase activity also has an off-target effect on MuSCs, which warrants the use of Tx-treated Pax7Cre^ERT2^ mice as experimental controls in future studies, and demand caution in interpreting data using other controls in previous studies.

## SUPPLEMENTAL FIGURES

**Supplemental figure 1.**
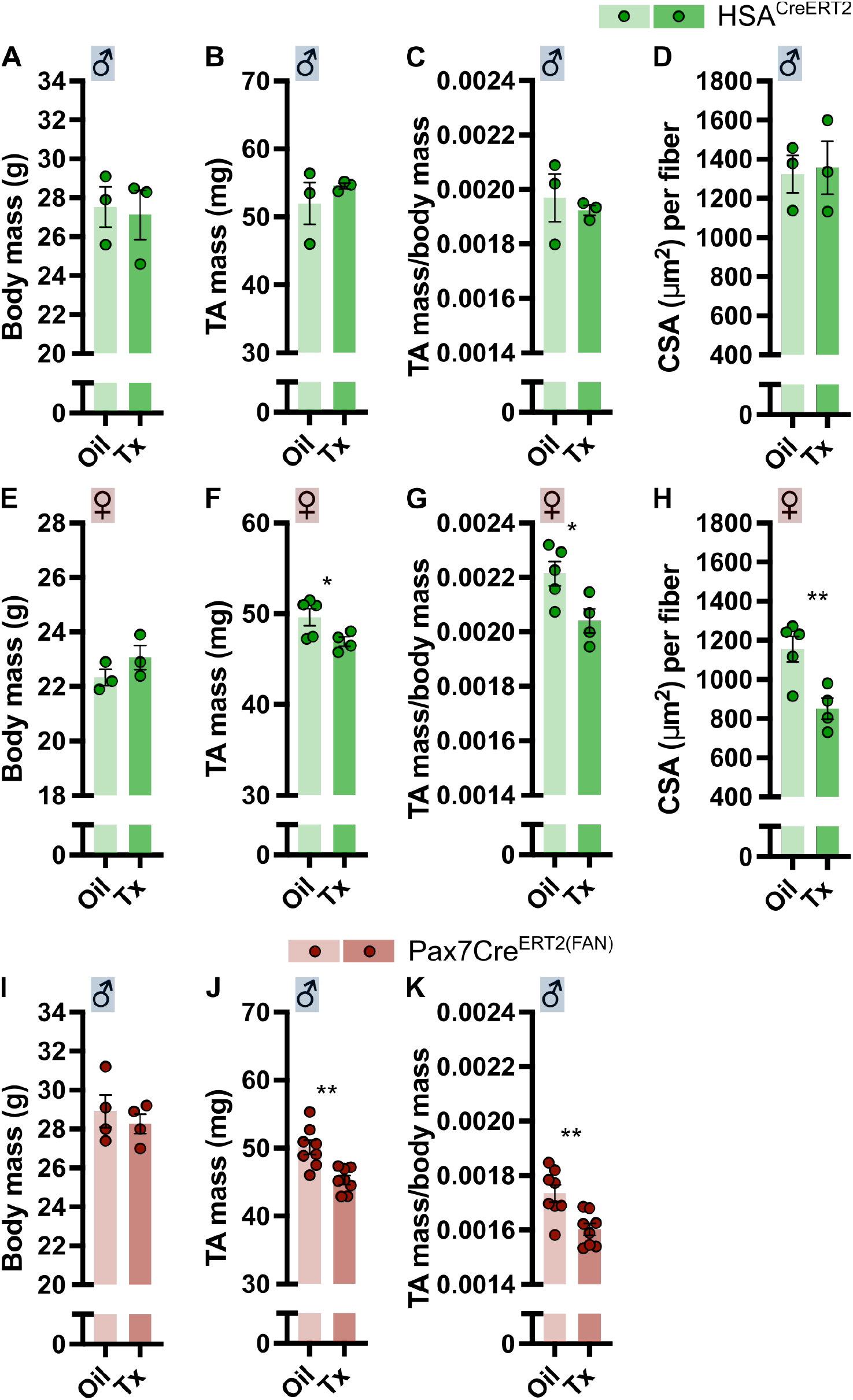
Skeletal muscle regeneration after Tamoxifen treatment at 14 days post injury in HSACre^ERT2^ mice, and after washout in Pax7Cre^ERT2^ mice. Male HSACre^ERT2^ mice treated with Oil or Tamoxifen (Tx) (A-D). A) Body mass, B) *Tibialis Anterior* (TA) mass, C) TA mass *per* mg body mass, D) average fiber cross sectional area (CSA). Female HSACre^ERT2^ mice treated with Oil or Tx (E-H). E) Body mass, F) TA mass, G) TA mass *per* mg body mass, H) average fiber CSA. Skeletal muscle regeneration at 14 days post injury in male Pax7Cre^ERT2(FAN)^ mice subjected to 14 days of washout before initiation of injury (I-K). I) Body mass, J) TA mass, K) TA mass *per* mg body mass. Bars represent means +/- SEM. *p<0.05 and **p<0.01.

**Supplemental figure 2.**
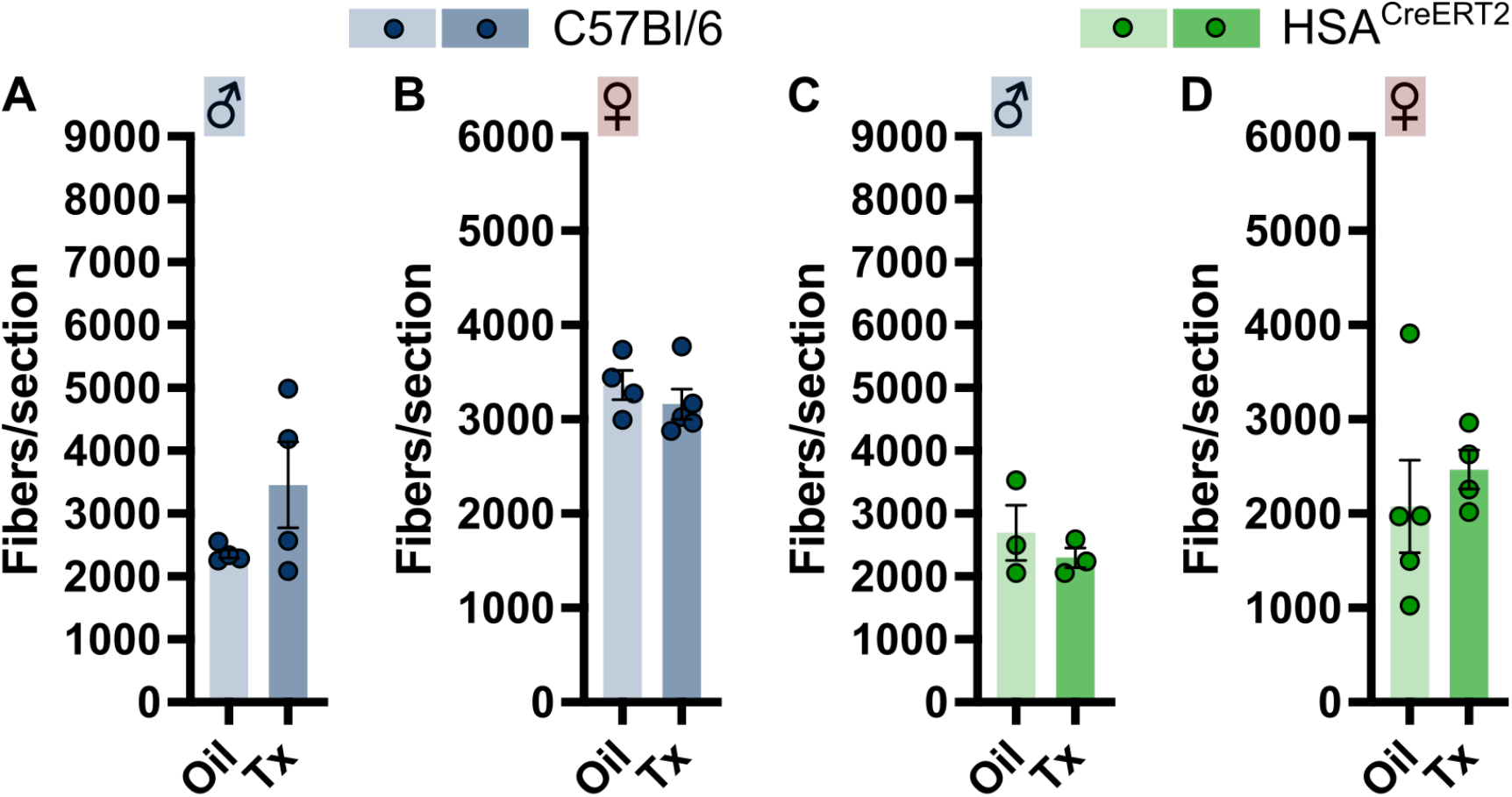
Number of fibers *per* section after Tamoxifen treatment at 14 days post injury in C57Bl/6 and HSACre^ERT2^ mice. A) number of fibers *per* section in male C57Bl/6 mice, B) number of fibers *per* section in female C57Bl/6 mice, C) number of fibers *per* section in male HSACre^ERT2^ mice, D) number of fibers *per* section in female HSACre^ERT2^ mice. Bars represent means +/- SEM.

**Supplemental figure 3.**
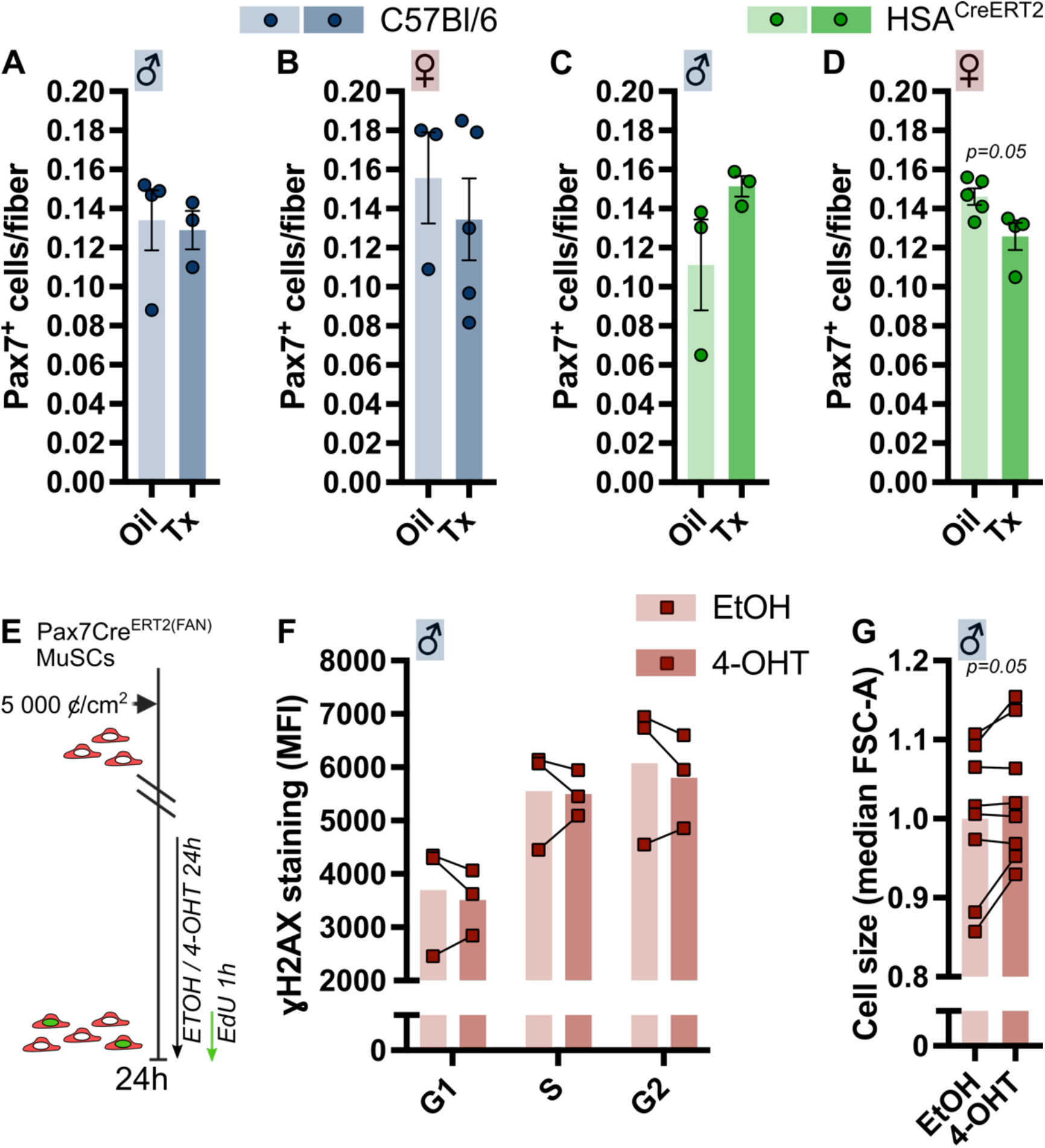
MuSC number and function after Cre activation. A) number of Pax7^+^ MuSCs *per* fiber at 14 days post injury (d.p.i.) in male C57Bl/6 mice, B) number of Pax7^+^ MuSCs *per* fiber at 14 d.p.i. in female C57Bl/6 mice, C) number of Pax7^+^ MuSCs *per* fiber at 14 days post injury (d.p.i.) in male HSA^CreERT2^ mice, D) number of Pax7^+^ MuSCs *per* fiber at 14 d.p.i. in female HSA^CreERT2^ mice, E) schematic of *in vitro* experiments on cells derived from male Pax7Cre^ERT2(FAN)^ mice in panel F-G, F) median fluorescence intensity of ɣH2AX staining *per* phase of the cell cycle by flow cytometry, G) cell size indicated as median FSC-A by flow cytometry. Bars represent means +/- SEM.

